# From tropical rainforests to scorching deserts: biogeography shapes climate-change plasticity in Australian rainbowfishes

**DOI:** 10.64898/2026.10.11.758404

**Authors:** Luciano B. Beheregaray, Katie Gates, Jonathan Sandoval-Castillo, Chris J. Brauer, Louis Bernatchez, Peter J. Unmack

## Abstract

Thermal tolerance and the plasticity of regulatory mechanisms are likely to influence biodiversity persistence under global warming. Continental-scale assessments of physiological resilience and gene expression plasticity within a closely related clade are rare, yet they could reveal how bioregional climatic history shapes vulnerability to climate warming. We used a comparative approach across five climatic ecotypes of Australian rainbowfishes (tropical rainforest, tropical savannah, subtropical, desert, and temperate) to investigate variation in thermal tolerance and gene expression plasticity in response to warming. We reconstructed phylogenetic relationships among these ecotypes, estimated their ecological niches and environmental warming tolerance, and experimentally measured CT_MAX_ and global gene expression responses to a projected summer heat event. Closely related ecotypes showed divergent bioclimatic niches, contrasting sensitivities to warming, and extensive but idiosyncratic expression responses to heat stress. Plastic responses to warming, including in key heat stress genes, were positively associated with thermal tolerance across ecotypes. The tropical savannah and subtropical ecotypes, both from environmentally variable bioregions, showed extensive transcriptional responses to warming and high thermal tolerance. The desert ecotype was intermediate for both traits, whereas the temperate ecotype showed the most limited transcriptional response and lowest thermal tolerance. In contrast, the tropical rainforest ecotype showed limited plasticity, potentially reflecting local adaptation to milder, more stable conditions and more specialised thermal responses. Our findings support the hypothesis that bioregional climatic history shapes broad patterns of climate resilience, including thermal tolerance and transcriptional plasticity, and may influence which combinations of traits are favoured under future climates.

## Introduction

Climatic bioregions provide a useful framework for understanding geographic variation in adaptation and resilience. Differences in climatic regimes and biotic and abiotic conditions have shaped physiological limits and adaptive traits, contributing to geographic variation in ecological niches, range sizes, and climate resilience (Bozinovic et al. 2011; Sunday et al. 2012; Stuart-Smith et al. 2017). As climates warm, organisms may increasingly face temperatures beyond those they have historically experienced, with their responses influenced not only by how much temperatures increase, but also by their existing physiological traits, phenotypic plasticity, and evolutionary capacity (Stillman 2003; Hoffmann and Sgrò 2011; Catullo et al. 2015). Because these attributes reflect the environments in which organisms evolved, thermal resilience is expected to vary geographically, making it important to understand how thermal limits vary across bioregions and the mechanisms influencing divergent responses (Thomas et al. 2004; Tewksbury et al. 2008; Pörtner et al. 2006; Bennett et al. 2021). Upper thermal limits alone however provide an incomplete measure of vulnerability because risk also depends on the temperatures that organisms already experience. Environmental warming tolerance, defined as the difference between an organism’s critical thermal maximum (CT_MAX_) and its environmental temperature, provides an estimate of the remaining thermal buffer before physiological limits are reached (Deutsch et al. 2008). Ecotypes with relatively high CT_MAX_ may remain vulnerable if they occupy environments where temperatures are already close to those limits. Ecological niche models provide a complementary means of characterising the climatic environments occupied by different ecotypes and testing whether their climatic associations are equivalent. Quantifying niche differentiation and overlap therefore provides the geographic context needed to interpret differences in environmental exposure, physiological tolerance and regulatory responses to warming.

Organisms from climatically stable environments may evolve specialised thermal physiologies, potentially at the cost of tolerance breadth and flexibility relative to those from more variable environments (Janzen 1967; Payne and Smith 2017; Bennett et al. 2021). This may be particularly important for tropical species, which often operate closer to their thermal limits and may have reduced acclimation capacity and fewer opportunities for behavioural avoidance than higher-latitude species (Ghalambor et al. 2006; Deutsch et al. 2008; Tewksbury et al. 2008; Huey et al. 2009; Pottier et al. 2025). Further variation likely exists among tropical regions, for example across elevation gradients or between rainforest and savanna bioregions, but the mechanisms underlying thermal resilience at this finer spatial scale remain poorly understood (Moritz et al. 2012; Polato et al. 2018; Gates et al. 2026; Holzmann et al. 2026). Much of this variation stems from the pervasive influence of temperature on ectotherm biology: body temperature affects almost all major components of physiology and behaviour (Angilletta et al. 2002; Hochachka and Somero 2002; Huey et al. 2009), and thermal tolerance itself can be shaped by a range of extraneous environmental factors (Beitinger 1990; Pörtner et al. 2017). Climate change compounds this complexity by increasing both the average and the variability of environmental temperatures (IPCC 2023), each of which may select for different thermal response traits (Gilchrist 1995; Angilletta Jr 2009). One key mechanism for responding to warming is phenotypic plasticity: the capacity of a single genotype to produce multiple phenotypes depending on environmental exposure (Scheiner 1993; Pigliucci 2001; Kelly et al. 2011; Sultan 2021). By broadening individuals’ thermal performance capacities, plasticity can buffer short-term stress while also facilitating longer-term adaptation to novel conditions (Pfennig et al. 2010; Wund 2012; Logan and Cox 2020; Bailey et al. 2021). It may effectively “buy time” until genomic adaptation occurs (Diamond and Martin 2021), or contribute to evolutionary feedback loops that accelerate adaptive evolution (Fusco and Minelli 2010; Bailey et al. 2021). This may be especially important for dispersal-restricted taxa (e.g., freshwater obligates), which have limited ability to relocate and receive little inflow of novel genetic variation (Muñoz et al. 2016; Brauer et al. 2017).

Plasticity is itself a heritable trait subject to selection (McCairns and Bernatchez 2012; McCairns et al. 2016; Goldstein and Ehrenreich 2021), and because its fitness costs and benefits vary across taxa and environments (DeWitt et al. 1998; Snell-Rood and Ehlman 2021), adaptive plasticity, like genetic variation, can differ geographically (Richards et al. 2006; Crispo 2008). Selection for plasticity is predicted across many scenarios, especially where individuals face variable environmental extremes over time or space (Baythavong 2011; Snell-Rood and Ehlman 2021). Although high plasticity can sometimes inhibit genetic adaptation (Oostra et al. 2018; Fox et al. 2019), it more often aligns with directional selection, providing additive or facultative benefits (Gonzalo Turpin and Hazard 2009; Jasienski 2009); fitness costs are reported less often, though formerly plastic traits may be lost through genetic assimilation in stable environments favouring specialisation (Kelly 2019; Scheiner and Levis 2021). Gene expression modification is among the most important mechanisms underlying plasticity, enabling acclimation across ambient conditions (Schlichting and Smith 2002; López-Maury et al. 2008; Logan and Cox 2020; Rivera et al. 2021) and revealing the pathways regulating environmental responses and likely targets of selection under long-term change (Alvarez et al. 2015; Gerken et al. 2015; Logan and Cox 2020). Comparing gene expression across populations or gradients can further reveal local reaction norms, which, alongside physiological indicators such as tolerance, help elucidate patterns of adaptive plasticity (McCairns and Bernatchez 2010; Whitehead 2012; Rivera et al. 2021).

Rainbowfishes (*Melanotaenia* spp.) are an emerging system for studying climatic resilience and adaptation. They represent the most diverse monophyletic group of freshwater fishes in Australia and New Guinea, occupying a broad range of aquatic habitats across these regions (Unmack et al. 2013). Being non-migratory, their population genomic diversity is often hierarchically structured at the scale of the catchment — a fundamental climatic unit of the natural landscape (Table 1 and Figure 1). Across their ranges, populations show differences in genotype–environment associations (GEAs) and local adaptation linked to hydroclimatic selection and to phenotypic traits known to affect fitness (Brauer et al. 2018; Smith et al. 2020; Attard et al. 2022; Brauer et al. 2023; Gates et al. 2023; Gates et al. 2026), making rainbowfishes a powerful field model for investigating climatic adaptive evolution at broad geographic scales. They are equally valuable as an experimental model, being small, sexually dimorphic, easy to maintain and breed, and having a short generation time. Experimental exposure to projected future climates has revealed predictable gene-expression plasticity in response to temperature stress for a subtropical species (Smith et al. 2013). Transgenerational experiments have provided pedigree-based evidence that key gene expression responses to projected summer temperatures are heritable (McCairns et al. 2016), with molecular pathways of candidate genes co-occurring with genes inferred to be under climate-mediated selection in wild populations (McCairns et al. 2016; Smith et al. 2020). Comparative climate-change experiments have revealed geographic variation in the genetic basis of adaptive plasticity in rainbowfishes: a subtropical species showed greater upper thermal tolerance and more responsive genes under thermal selection than temperate and desert species (Sandoval-Castillo et al. 2020). Together, these rainbowfish findings suggest that climate resilience may be shaped by the biogeographic context in which populations evolved (Armarego-Marriott 2021). Yet comparative studies of thermal tolerance and gene-expression plasticity remain uncommon (Levine et al. 2011; Logan and Buckley 2015; King et al. 2018), particularly in the tropics, a biodiversity-rich but poorly studied region that is highly vulnerable to warming (Beheregaray et al. 2015; Polato et al. 2018; Tewksbury et al. 2008). In tropical organisms, warmer and more variable environments may favour greater thermal response capacity, whereas historically stable environments may promote greater specialisation or assimilation of these responses (Gilchrist 1995; Tewksbury et al. 2008; Kelly 2019; Holzmann et al. 2026).

**Figure 1.**
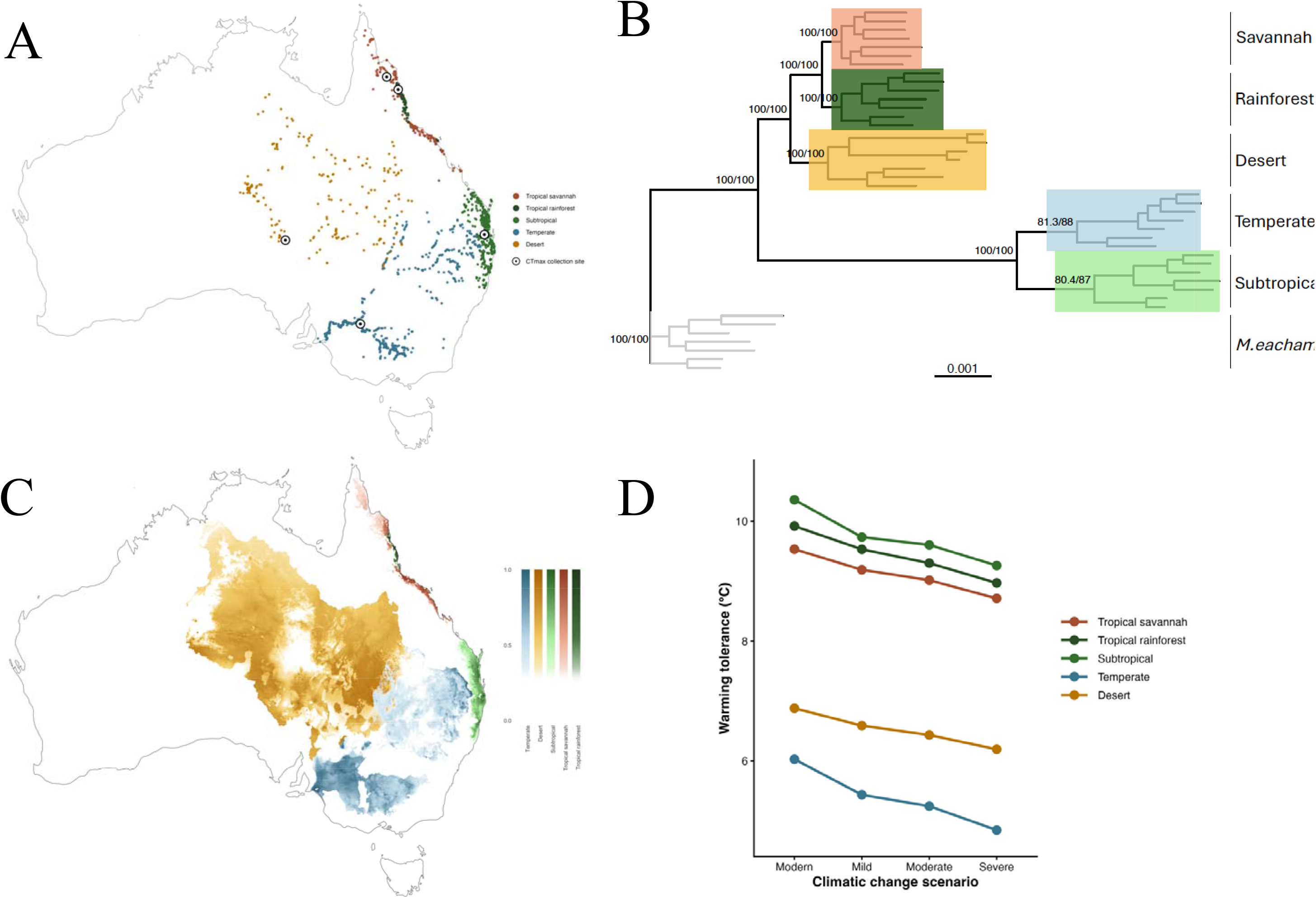
**(A)** Taxonomically validated occurrence records for the five *Melanotaenia* ecotypes based on 8,623 occurrence records from Atlas of Living Australia (ALA 2026). **(B)** Maximum likelihood tree built with IQ-TREE v3 showing phylogenetic relationships among *Melanotaenia* ecotypes based on ∼0.72 million aligned sites; the tree was rooted with *M. eachamensis*. **(C)** Ecological niche modelling for the five ecotypes based on 1,488 retained occurrence cells and four bioclimatic variables. Color tone indicates habitat suitability of each ecotype. **(D)** Physiological sensitivity of ecotypes across their ranges estimated by the environmental warming tolerance quantity for modern days and three shared socioeconomic pathways to represent mild (SSP126), moderate (SSP245) and severe (SSP585) climate change scenarios.

**Table 1.**
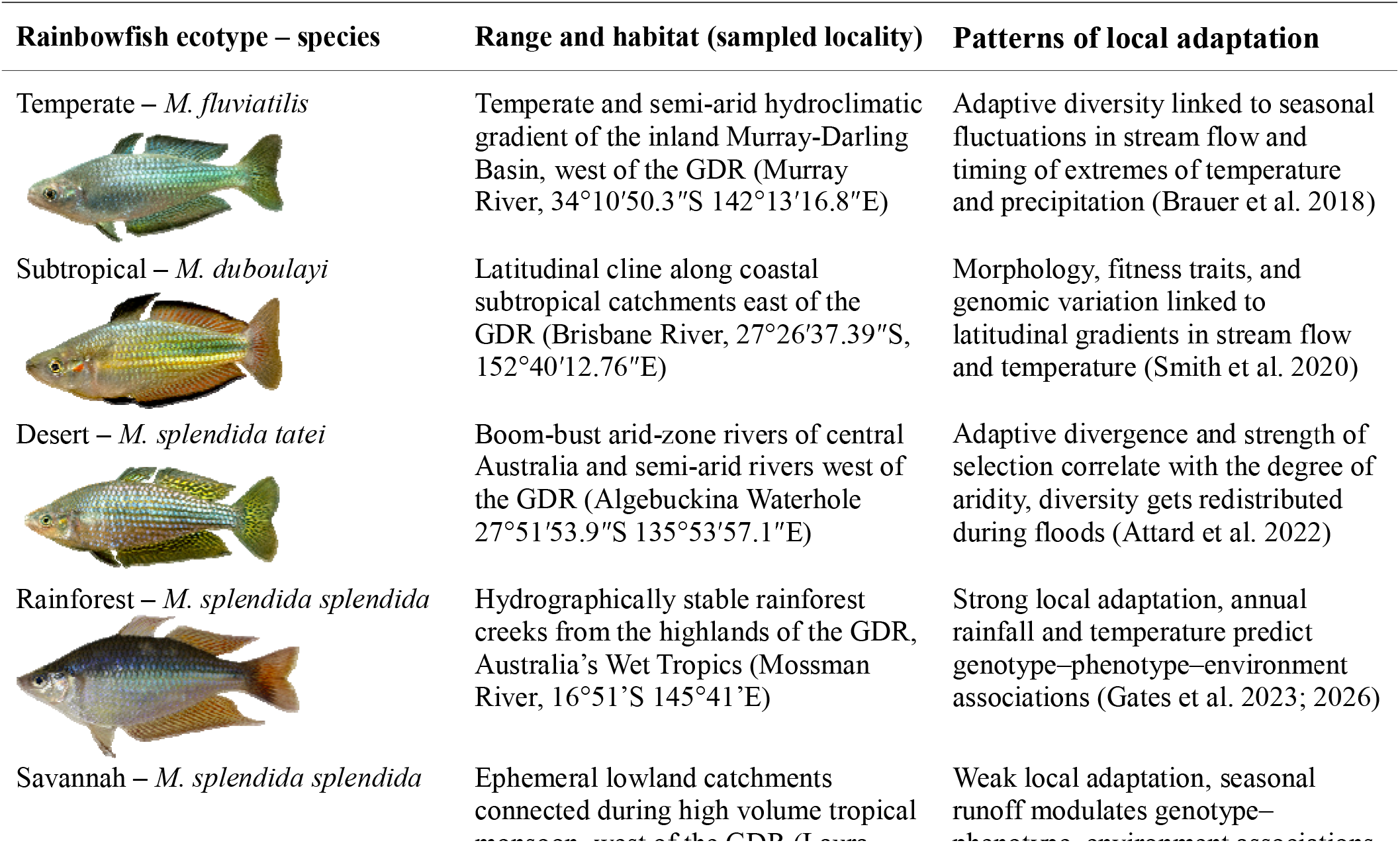
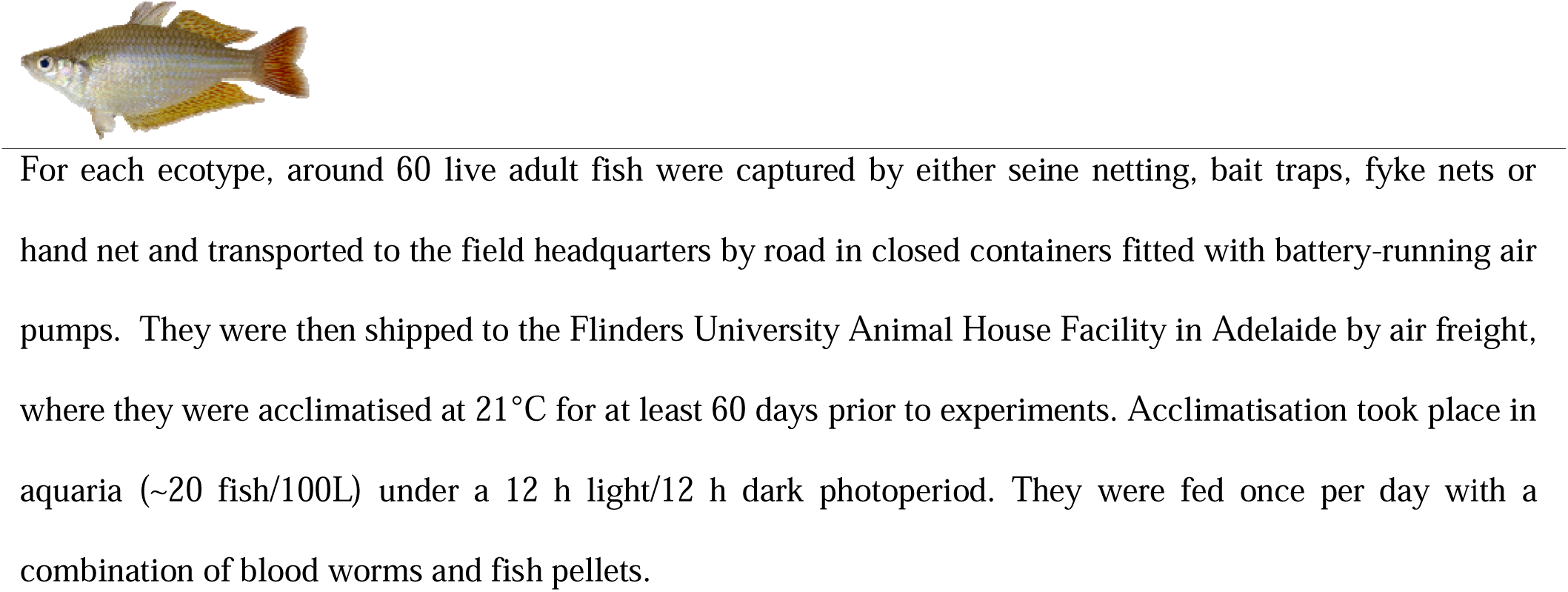
The climatic ecotypes of rainbowfishes (*Melanotaenia*), their range and habitat, sampled locality for experiments and environmental predictors of adaptation based on range-wide studies of landscape genomics.

Here we used an integrative comparative approach to study the biogeographic basis of physiological resilience and gene expression responses to an extreme heat event. We targeted five locally abundant rainbowfish taxa representing distinct “climatic ecotypes” (sensu Engelhard et al. 2010), each endemic to a distinct major contemporary climatic bioregion of Australia: tropical rainforest, tropical savannah, subtropics, desert, and temperate (Figure 1, Table 1). We first used genome-wide data to reconstruct phylogenetic relationships among these ecotypes and test whether they are closely related. We then estimated each ecotype’s ecological niche and their environmental warming tolerance to delineate climatic envelopes, test for niche identity, and assess physiological sensitivity across their ranges. Finally, we experimentally measured CT_MAX_ and global gene expression profiles in response to projected climate warming for each ecotype, and created a protein interaction network to assess the functional relevance of thermally responsive genes. Interpretation of these spatial and experimental results was facilitated by existing knowledge of range-wide hydroclimatic adaptation for the five ecotypes (summarised in Table 1), illustrating the benefits of combining landscape genomics with experimental studies of climatic resilience.

Our study builds on comparative work of rainbowfishes at higher latitudes (Sandoval-Castillo et al. 2020) to deliver a rare continent-wide assessment of climate-change plasticity within a closely related clade. It incorporates data from two biodiversity-rich but poorly studied tropical ecosystems: the relatively stable and topographically complex rainforest and the more environmentally variable and spatially connected savannah. Our central premise is that bioregional climatic history shapes broad patterns of climate resilience, including thermal tolerance and transcriptional plasticity, and influences the combinations of traits likely to be favoured under future climates (Janzen 1967; Bennett et al. 2021; Hawkins et al. 2003). We expected bioregion-specific responses, including greater heat tolerance in ecotypes that evolved at lower latitudes (Deutsch et al. 2008; Sunday et al. 2011, 2019) and idiosyncratic, rather than repeatable (deMayo and Ragland 2025) transcriptional responses to heat stress among ecotypes. We further hypothesised that transcriptional plasticity contributes to thermal resilience (Ghalambor et al. 2015), predicting a broad continental-wide positive relationship between upper thermal limits and the number of genes differentially expressed under warming. Conversely, we expected less flexible transcriptional responses, consistent with genetic assimilation or local adaptation, in ecotypes inhabiting less connected metapopulations that evolved in naturally fragmented and temporally stable environments.

## Methods

### Ecotype characterisation and sampling for experiments

The higher latitude ecotypes targeted here include the crimson spotted rainbowfish (*M. duboulayi*), a species found in subtropical coastal catchments east of Australia’s Great Dividing Range (GDR); the Murray River rainbowfish (*M. fluviatilis*), a species found in temperate catchments of the inland Murray-Darling Basin west of the GDR; and the desert rainbowfish (*M. splendida tatei*), a species found in arid and semiarid catchments of central Australia. The targeted tropical ecotypes are from the eastern rainbowfish (*M. splendida splendida*), a species endemic to Queensland that shows extensive variation in colour and morphology across their range, which includes both rainforest and savannah ecoregions (Table 1, Figure 1 A). The rainforest ecotype is found in densely packed, and mostly perennial drainage networks in the rugged terrain of the GDR (Nott 2005, Pearson et al. 2015). In contrast, the savannah ecotype is found in often ephemeral and interconnected lowland drainage systems west of the GDR. The populations sampled for the two tropical ecotypes are geographically isolated from each other and highly divergent morphologically and genomically (Gates et al. 2026).

### Phylogenetics

Phylogenetic relationships among Melanotaenia ecotypes were inferred using a partitioned maximum-likelihood approach implemented in IQ-TREE v3 (Wong et al. 2026). The tree was based on 2,777 AD loci comprising approximately 0.72 million aligned sites, generated using ddRAD (double-digest restriction-site-associated DNA sequencing) as part of five range-wide landscape genomic studies summarised in Table 1. Following quality filtering and trimming, sequences were mapped to a new chromosome-level reference genome assembly for M. s. splendida. Regions covered by at least 80% of individuals, with a minimum sequencing depth of four reads, were retained. The concatenated alignment was partitioned by RAD locus, and the optimal substitution model for each partition was selected using ModelFinder in IQ-TREE. Node support was assessed using 1,000 ultrafast bootstrap and 1,000 SH-aLRT replicates. To account for intraspecific geographic variation, samples from the most genomically divergent populations identified in previous studies were included. The tree was rooted with M. eachamensis, a closely related species from the Atherton Tablelands in Australia’s Wet Tropics (Unmack et al. 2013). All taxa included in the analysis belong to the relatively young ‘australis’ clade, which forms part of the broader and widely distributed ‘southern lineage’ of Melanotaenia rainbowfishes (Unmack et al. 2013).

### Ecological niche modelling and environmental warming tolerance

A total of 8,623 occurrence records were obtained from the Atlas of Living Australia (ALA 2026) and retaining one occurrence per taxon per 2.5-arc-minute climate cell, 1,488 occurrence cells remained: 188 tropical-savannah *splendida*, 87 tropical-rainforest *splendida*, 446 *duboulayi*, 557 *fluviatilis*, and 210 *tatei*. The *splendida* occurrences were classified using the 1991–2020 Köppen–Geiger climate classification (Beck et al. 2018), with Af and Am climates defining tropical rainforest and Aw tropical savannah. Baseline WorldClim 2.1 bioclimatic variables (1970–2000; Fick et al. 2017) were aggregated to a 2.5-arc-minute grid. BIO5 and BIO6 were retained a priori because of their relevance to thermal adaptation, while remaining predictors were screened using Pearson correlation (≤0.8). Presence-background niche models were fitted with the maxnet v2.0.5.2 R package (Phillips et al. 2017), using taxon-specific river regions for calibration and up to 3,000 mainland Australian background cells. Nine combinations of feature classes and regularization multipliers were evaluated using four-fold spatial cross-validation in the ENMeval v0.1.4 R package (Kass et al. 2021). Models with mean withheld omission rates >0.15 were excluded, and settings were then selected based on withheld AUC, model complexity and regularization. For niche comparisons, models were standardised to BIO5, BIO6, BIO12 and BIO15. Pairwise niche identity was tested using ENMTools with Maxent 3.4.3, using Schoener’s D and Hellinger-based I. Significance was assessed from 999 randomisations, with Holm correction applied separately to each metric.

Environmental warming tolerance (Deutsch et al. 2008) was calculated for each ecotype as mean experimental CT_MAX_ (details below) minus the mean estimated warmest month stream temperature across retained occurrence cells. For each occurrence cell, the maximum temperature of the warmest month was identified from 12 monthly maximum air temperature layers and converted to stream temperatures using the Punzet et al. (2012) global regression. Modern conditions were represented by the WorldClim 2.1 data for 1979-2000. Future estimates were based on BCC-CSM2-MR projections for 2061–2080 under three shared socioeconomic pathways to represent mild (SSP126), moderate (SSP245) and severe (SSP585) climate change scenarios.

### Inferring thermal tolerance

We determined the upper thermal limit for organised locomotory activity (CT_MAX_) for each ecotype following the methods of Becker and Genoway (1979), which have previously been applied to rainbowfishes (Smith et al. 2013; McCairns et al. 2016; Sandoval-Castillo et al. 2020). We used 10 similarly sized adult females from each of the five ecotypes acclimated at 21°C, for a total of 50 individuals. Each fish was placed individually in a 5-L glass beaker containing 3.5 L of aquarium water, with the beaker partially submerged in a digital water bath (SWBD, Stuart®). Water temperature in the beaker was increased by approximately 1°C every three minutes (0.33°C min ¹) until CT_MAX_ was reached (Becker and Genoway 1979). CT_MAX_ was defined as the loss of righting response, whereby a fish was unable to maintain its normal dorsal–ventral orientation for at least one minute (Monirian et al. 2010). This endpoint represents a level of thermal stress at which an animal is no longer able to maintain essential functions or escape from the source of stress (Becker and Genoway 1979; Monirian et al. 2010). Once the loss of righting response was observed, the time and temperature were recorded, and the fish was transferred to a recovery tank containing aerated water at its acclimation temperature. Fish were monitored for 24 h after the experiment for behavioural abnormalities or mortality. CT_MAX_ for each ecotype was calculated as the mean of the 10 independent replicate measurements. We compared CT_MAX_ among ecotypes using analysis of variance (ANOVA).

### Trial of an extreme heat event

We simulated an extreme and prolonged heat event likely to trigger overheating that is directly comparable with trials used in previous experiments of Australian rainbowfishes (Smith et al. 2013; Sandoval-Castillo et al. 2020). The treatment temperature (33°C) represents a projected summer temperature for eastern Australia in 2070 under a high-emission scenario (SSP585). Although the growth rate of global carbon emissions is slowing, this high-emission scenario remains relevant given projected warming by 2100, with its associated temperatures considered a reasonable match for mid-century warming under current and stated policies (Schwalm et al. 2020). To assess short-term responses to contemporary and projected summer temperatures, similarly-sized adult males from each ecotype were randomly assigned to either a treatment (33°C) or control (21°C) group. Following the protocol of Smith et al. (2013), temperatures in treatment tanks were increased by 2°C per day over six days until reaching the target temperature of 33°C, which was then maintained for 14 days. Control groups were maintained at 21°C throughout the experiment. Between 9:00 am and 11:00 am, fish from both treatment and control groups were euthanised using an overdose of anaesthetic (AQUI-S®, 175 mg/L for 20 min), following the Australian Code of Practice for the Care and Use of Animals for Scientific Purposes (2013). Immediately after death, individuals were dissected and the liver extracted. We focused on the liver because of its established role in metabolic conditioning during heat-stress responses and relatively homogeneous distribution of cell types (Rabergh et al. 2000; Smith et al. 2013). Liver tissue was placed in RNAlater (Ambion) and incubated at 4°C for 12 h before being transferred to −80°C.

### RNA sequencing (RNA-seq) to characterise global gene expression

Total RNA was extracted from liver samples of 60 rainbowfish (six individuals per experimental group; 12 per ecotype) using the MagMAX™-96 Total RNA Isolation Kit (Ambion), including a DNase treatment. RNA quantity and quality were assessed on a Bioanalyzer 2100 (Agilent Technologies). Libraries were prepared with the TruSeq RNA™ Sample Preparation LT Protocol (Illumina): poly-A mRNA was isolated, fragmented, and reverse-transcribed to cDNA, which was end-repaired, 3′-adenylated, and ligated to unique indexed adapters (MID tags 2, 4-7, 12-16, 18, 19). The bead ratio during PCR clean-up was adjusted to exclude short fragments (<200 bp). Libraries were normalised, pooled in groups of 12 fish per lane (Smith et al. 2013), and sequenced on an Illumina HiSeq 2500 at Novogene (Hong Kong) to generate 100-bp paired-end reads

### Bioinformatics: quality trimming, genome-guided assembly, and quality assessment

Reads were quality-trimmed with TRIMMOMATIC v0.36 to remove adapters, low-quality bases (Q < 20), and reads shorter than 45 bp. Trimmed reads were mapped to the *Melanotaenia* transcriptome (Sandoval-Castillo et al. 2020), and the alignments were used to assemble a new and improved *Melanotaenia* transcriptome with TRINITY v2.5.1. Assembly quality was assessed from the percentage of raw reads retained, contig N50, and the percentage of full-length transcripts. Candidate protein-coding regions (ORFs ≥100 amino acids) were identified with TRANSDECODER v3.0, and transcripts sharing ≥80% sequence similarity were collapsed to the longest representative to produce a non-redundant set of unigenes (hereafter genes). To remove extraneous genes, we mapped the genes to the chromosome-level *M. s. splendida* reference genome and retained only the matching sequences. Genes were annotated by BLASTX against the UniProtKB Danio rerio database (e-value ≤ 1 × 10 6), and transcripts with ≥50% similarity to bacterial, fungal, or viral genes were removed.

### Differential expression and network analyses

Reads from each individual were mapped to the *M. s. splendida* predicted coding regions with BOWTIE2 v2.2.7, and gene-level abundance was estimated with RSEM v1.2.19. Counts were normalised across samples using the trimmed mean of M values (TMM), and differential expression (DE) was tested with DESeq2 v1.10.1. Following Sandoval-Castillo et al. (2020), DE genes were defined as those with a ≥ log2 fold-change at 5% FDR between any two ecotype–treatment groups (e.g. rainforest-experimental vs. rainforest-control, or rainforest-experiment vs. savannah-experiment). Only the DE genes between treatment (experimental and control) of each ecotype were considered. This addresses only activational plasticity, i.e. expression changes that are induced, and often reversed, within an individual’s lifetime (Snell-Rood and Ehlman 2021). Relationships between expression plasticity and upper thermal tolerance were tested by linear regression of the number of experiment-vs-control DE genes against thermal tolerance for each ecotype.

To explore system-level functions among thermally induced genes and identify those central to the response, protein interaction networks were built in CYTOSCAPE v3.7 (Rakshit et al. 2014) from thermally DE genes in the combined dataset. Edges linked genes with reported physical or functional interactions in zebrafish (*Danio rerio*) in the STRING database (Szklarczyk et al. 2016). Gene importance was evaluated by node degree, and genes with degrees ≥ the mean + 2 SD of the node degree distribution were classed as hubs (Rakshit et al. 2014; Sandoval-Castillo et al. 2020).

## Results

### Phylogeny

The maximum likelihood analysis of genome-wide data was consistent with the current taxonomy and supported a close evolutionary relationship among the five ecotypes (Figure 1 B). The tropical rainforest and savannah ecotypes (*M. splendida splendida*) clustered with the desert ecotype (*M. s. tatei*), forming the *splendida* clade, while the subtropical ecotype (*M. duboulayi*) was sister to the temperate ecotype (*M. fluviatilis*). Within each taxon and ecotype, all geographic population samples formed a well-supported monophyletic group.

### Ecological niche modelling and environmental warming tolerance

Model performance was strongest for the tropical savannah and rainforest groups, with mean spatial cross-validation AUCs of 0.842 (SD = 0.102; omission = 0.069) and 0.803 (SD = 0.119; omission = 0.125), respectively. Support was more modest for the subtropical (AUC = 0.609, SD = 0.069; omission = 0.095) and temperate (AUC = 0.621, SD = 0.123; omission = 0.245) models, while the desert model showed weak discrimination (AUC = 0.537, SD = 0.102; omission = 0.144) (Table S1). The savannah and rainforest ecotypes showed the greatest climatic niche overlap (Schoener’s D = 0.430; I = 0.736). Among the other ecotypes, overlap was highest between temperate and desert (D = 0.330; I = 0.632), followed by subtropical and temperate (D = 0.227; I = 0.470) and subtropical and desert (D = 0.147; I = 0.378). In contrast, comparisons between either tropical ecotype and the other species showed little overlap (all D < 0.10; I < 0.29) (Figure S1). Despite these differences in overlap, niche identity tests indicated significant differentiation for all ten pairwise comparisons: observed D and I values were lower than all 999 randomised values (raw P = 0.001; Holm-adjusted P = 0.010 for both metrics) (Table S2). Thus, all pairwise comparisons rejected climatic niche equivalence, while the magnitude of overlap indicated moderate similarity between the two tropical *splendida* ecotypes but limited overlap between tropical and non-tropical groups.

Modern warming tolerance was highest for the subtropical ecotype (10.36°C), followed closely by the rainforest (9.92°C) and savannah (9.53°C), and then by the desert (6.88°C) and temperate (6.03°C) ecotypes. Warming tolerance declined progressively under increasingly severe climate-change scenarios for all ecotypes (Fig 1D). Under SSP585, projected warming tolerance declined to 9.26°C for subtropical, 8.97°C for rainforest, 8.71°C for tropical savannah, 6.19°C for desert and 4.84°C for temperate rainbowfish. The temperate ecotype therefore had the lowest estimated warming tolerance under both modern and projected conditions and showed the largest decline under SSP585 (1.18°C), whereas the desert ecotype showed the smallest decline (0.68°C).

### Gene expression differences among temperature treatments and ecotypes

The *Melanotaenia* transcriptome assembly comprised 450,246 contigs and 315,268 ‘Trinity genes’, of which 64,694 were identified as ORFs (Table S3). A final subset of 36,536 ‘genes’ were found to be non-redundant across all ecotypes and used for subsequent DE analyses (Figure 2A). Of all genes, 78.8% (28,781) were expressed in all ecotypes, and 80% were expressed in at least four of the five ecotypes (Figure 2A). In marked contrast, none of the 520 genes that were DE between climate change treatment groups (Figure 2B) were shared across all ecotypes, and only six genes were DE in at least four ecotypes. Only 24 of these 520 genes were DE in the three *splendida* ecotypes, and 55 were DE in both the rainforest and savannah ecotypes. Considering the 5,000 most DE transcripts, responses were most highly correlated among individuals within ecotypes, followed by high correlation between control and experimental treatment groups within each ecotype (Figure 3A). These patterns indicate strong differential plasticity of expression across ecotypes induced by the projected heat event as well as phylogenetic signal in expression profiles. While thermal response was the most important factor driving pairwise gene expression differences among individuals (Figure 3A), lineage-specific responses drove expression differences within treatments (Figure 3B). The most flexible transcriptional response to warming was found for the savannah ecotype (205 DE genes), followed by the subtropical (149), the desert (137), the rainforest (111) and the temperate (64) ecotype (Figure 2B, Table S4).

**Figure 2.**
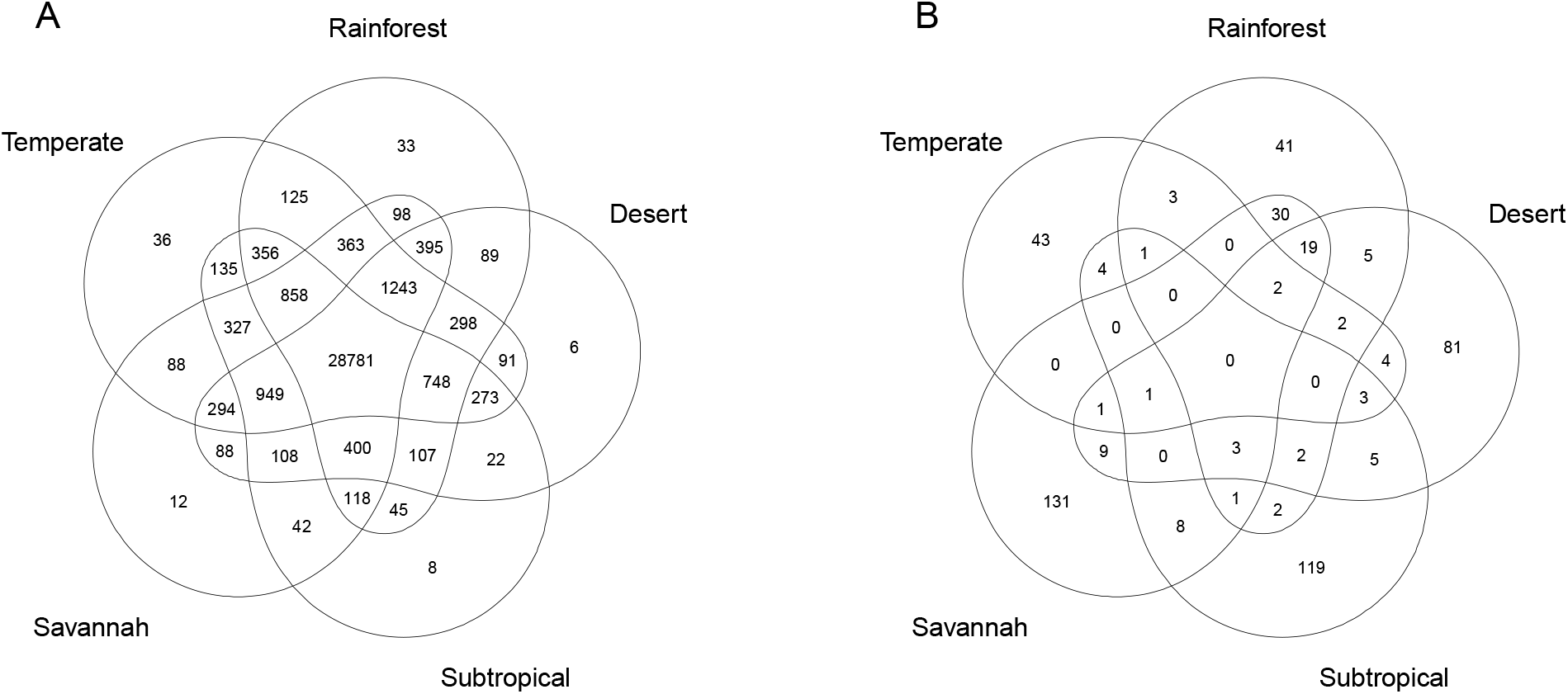
Venn diagram of global gene expression shared among the five *Melanotaenia* ecotypes, based on **(A)** all 36,537 identified *Melanotaenia* genes, and **(B)** genes differentially expressed between control (21°C) and projected 2070 summer treatment (33°C) groups (blank segments indicate no shared responses).

**Figure 3.**
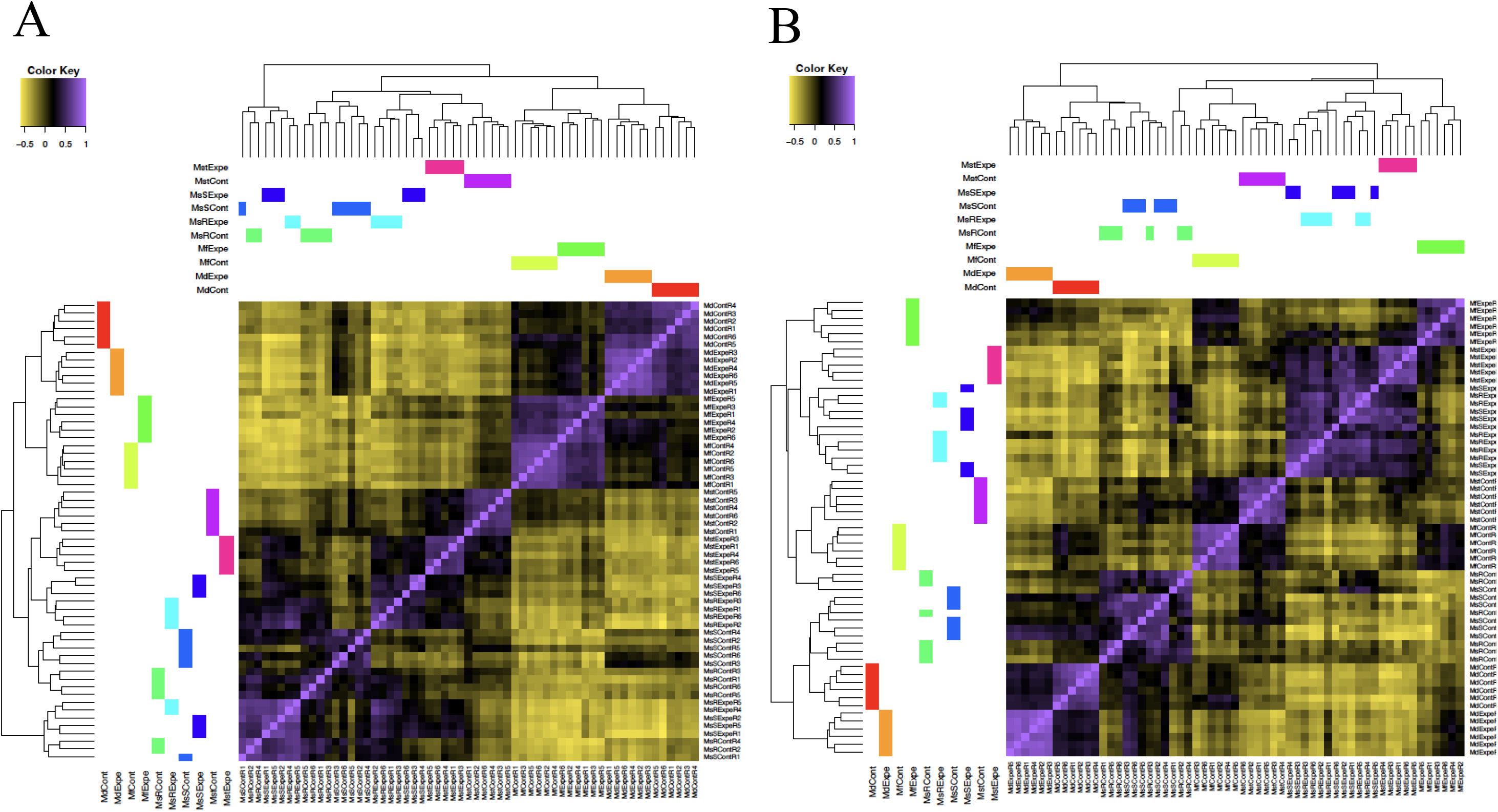
**(A)** Heatmap of the correlation matrix among the five *Melanotaenia* ecotypes in pairwise log_2_ gene expression profiles. This analysis was based on the 5,000 most DE transcripts. **(B)** Heatmap of the correlation matrix between treatments (control vs. experiment) in pairwise log_2_ gene expression profiles. This analysis was based on 520 DE genes identified between control and experiment. Coloured bars represent ecotypes; Expe = experimental climate change group, Cont = experimental control group.

### Upper thermal tolerance

At the individual level, CT_MAX_ ranged from 33.1°C for the temperate to 39.0°C for the savannah ecotype. On average, CT_MAX_ was higher in the savannah (38.4°C, SD = 0.3°C) and in the rainforest (38.2°C, SD = 0.5°C). The mean upper thermal tolerance of tropical *splendida* overall (38.3°C, SD = 0.4°C) was significantly higher (*P* < 0.05), than that of subtropical (38.0°C, SD = 0.4°C), desert (37.2°C, SD = 0.5°C), or temperate (34.9°C, SD = 1.2°C) rainbowfishes (Figure 4, Table S5, S6). Among the five ecotypes, a significant linear relationship was detected between CT_MAX_ and the number of genes DE between climate change treatments (*r* = 0.909, Figure 4).

**Figure 4.**
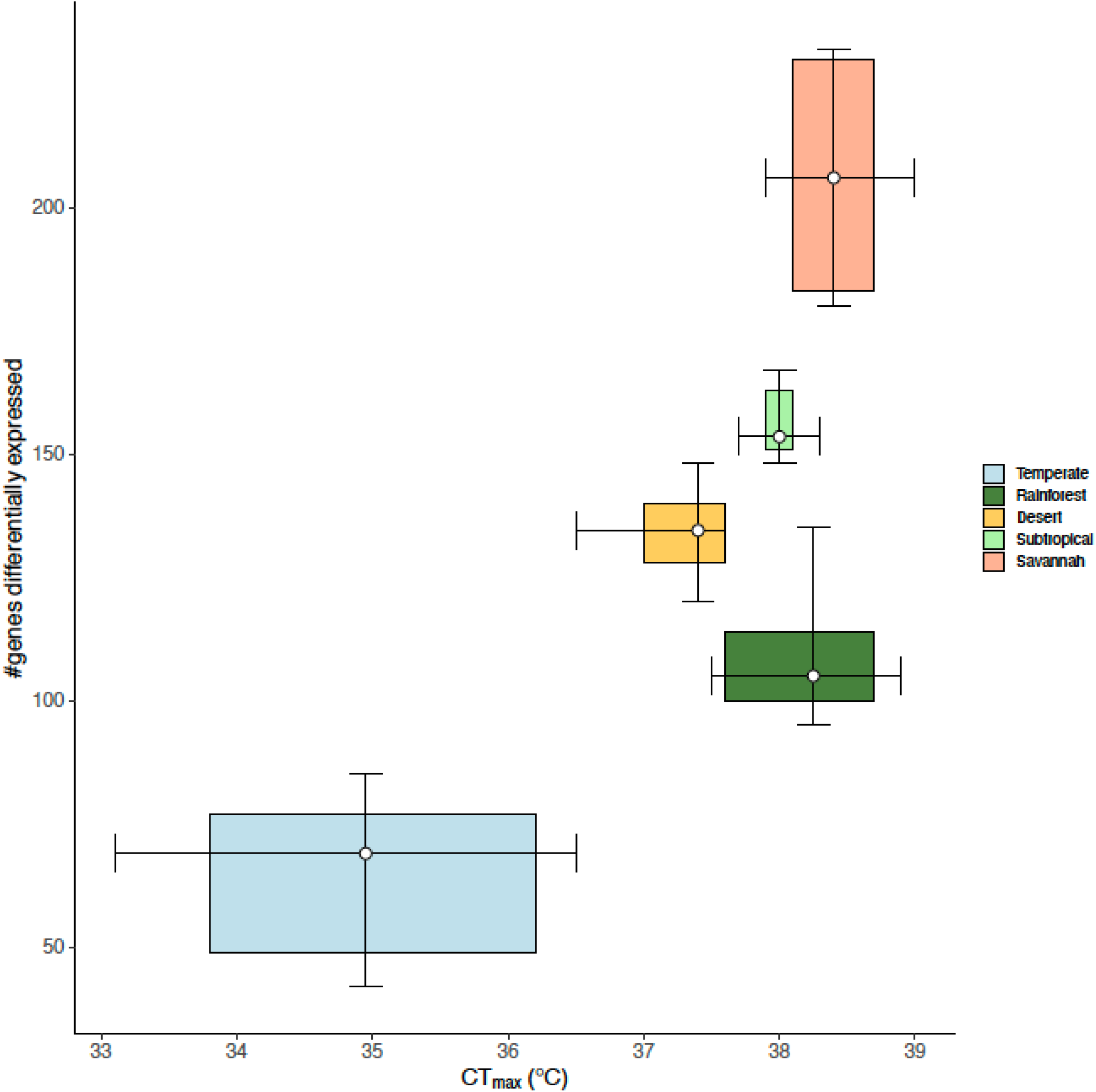
Associations between CT_MAX_ and the number of differentially expressed genes between control (21°C) and projected 2070 summer treatment (33°C) groups among five Australian *Melanotaenia* ecotypes (*r* = 0.909). The box plots display the upper and lower quartiles, whiskers represent 95th and 5th percentiles, and their intersections represent the median.

### Functional annotation and network analysis

Transcript annotation produced 26,919 protein hits, 25,169 (93.5%) of which were assigned to 317,948 gene ontology (GO) terms (Figure S2; Table S8). For the subset of genes DE under climate change treatments, we received 413 protein hits corresponding to 3,407 GO terms, for the rainforest these numbers were 103 protein hits and 936 GO terms, while for the savannah 176 protein hits with 1,250 GO terms. Protein networks for annotated DE genes were able to be constructed without requiring additional interactions (Figure 5). From the network analyses, we identified 13 hub genes for warming responses across all ecotypes, defined by high degrees of interaction within protein networks. Of these, nine were DE in savannah (AACS, ACLYA, ACLYB, CTPS1A, CYP51, DHCR24, FDFT1, HMGCS1 and LSS), and one was thermally induced in both ecotypes (ACLYA).

**Figure 5.**
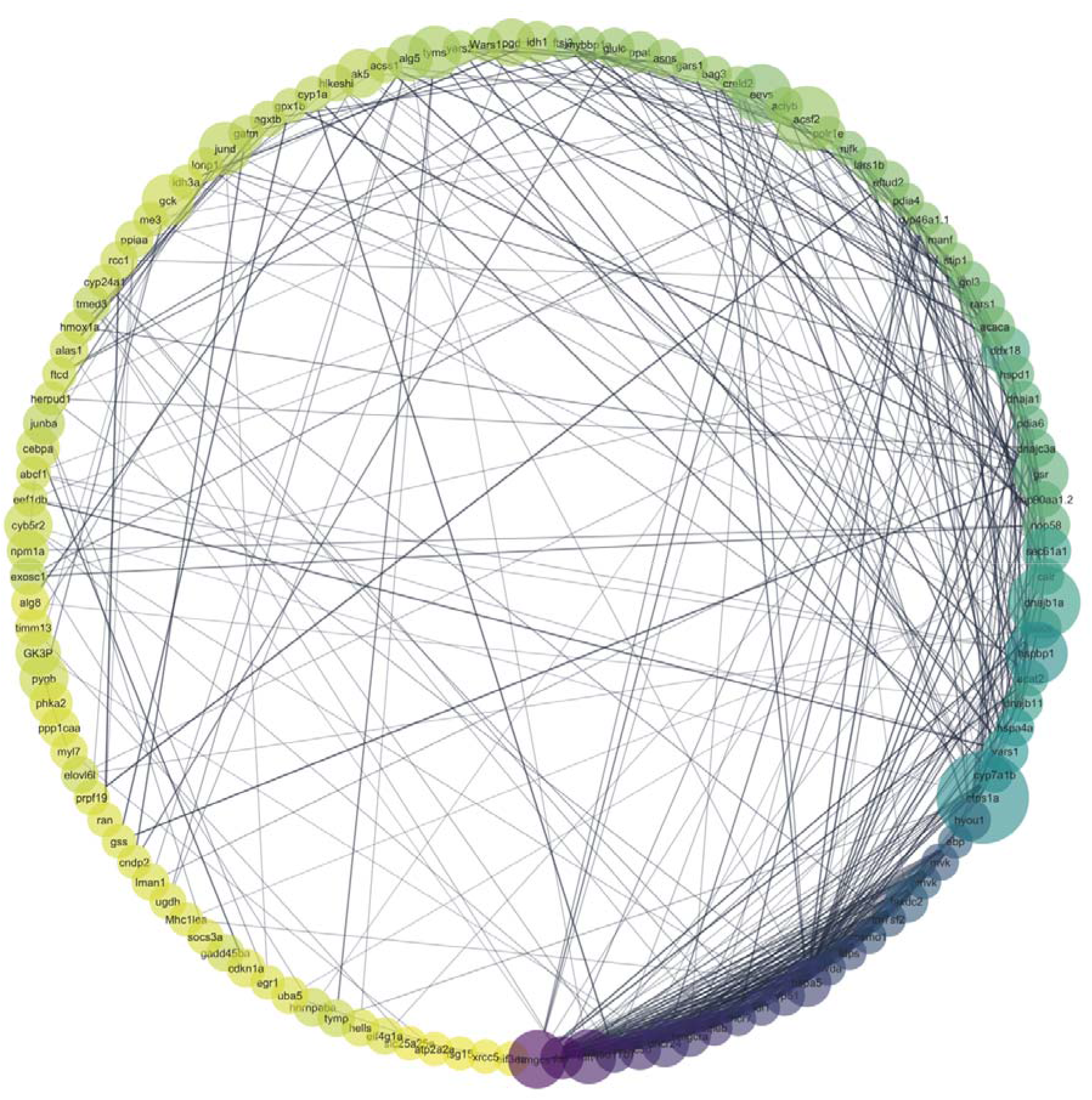
Protein interaction network for climate warming responses in *Melanotaenia* ecotypes based on 520 genes differentially expressed between control (21°C) and projected 2070 summer treatment (33°C) groups. Node sizes are proportional to centrality in the network (betweenness centrality), while shading indicates the relative number of direct interaction (neighbourhood connectivity; green = fewer interactions, orange = more interactions).

## Discussion

We implemented a comparative continent-wide study of five rainbowfish ecotypes to assess how bioregional climatic history influences plasticity and resilience to climate change. Using a clade distributed from tropical to temperate regions of mainland Australia, we found that closely related ecotypes nonetheless show divergent bioclimatic niches, along with differing range-wide sensitivities to environmental warming and to a projected heat event. Although they have very similar transcriptomes (e.g. 28,864 genes were shared across all ecotypes) and mounted extensive expression responses to warming, their plasticity was highly idiosyncratic (e.g. no genes were shared across all ecotypes in the climate experiment). Plastic responses and estimated upper thermal tolerance varied across bioregions, supporting our hypothesis that transcriptional plasticity facilitates thermal resilience in Australian rainbowfishes. Indeed, there was a strong positive relationship between climate-change-induced expression responses (including in key heat stress genes) and thermal tolerance across the five ecotypes.

### Contrasting plasticity and resilience to climate change within a tropical species

Although climate warming is often expected to have weaker impacts in the tropics than at higher latitudes, mounting evidence suggests that tropical ectotherms are highly susceptible to rising temperatures, including overheating risk, and that they are approaching the evolutionary limits of their thermal tolerance (Tewksbury et al. 2008; Pottier et al. 2025; Holzmann et al. 2026). Thermal resilience in tropical organisms has received increasing attention, particularly along elevation gradients (Polato et al. 2018; Holzmann et al. 2026), but comparative studies of gene expression plasticity across tropical climatic bioregions remain rare.

We found that transcriptional plasticity to projected warming differed markedly between two tropical ecotypes of the same species, each occurring in a different climatic bioregion. In the tropical rainforest ecotype, 111 genes responded to the simulated heat event, compared with 205 differentially expressed (DE) genes in the savannah ecotype. These two ecotypes shared only 10.6% of all 520 DE genes identified in the climate-change experiment. This strong divergence in thermally induced expression contrasts with their highly similar constitutive gene expression, with 32,256 sequenced genes (88.3% of the total) detected in both ecotypes. This suggests that constitutive expression may be more conserved between the rainforest and savannah ecotypes, whereas selective or demographic influences may have contributed to their distinct thermally induced expression profiles (Whitehead and Crawford 2006). Selection may influence plasticity through changes in both the number and identity of expressed genes and the magnitude of changes in gene expression (Schlichting and Smith 2002; Komoroske et al. 2021). Our previous work showed that a subtropical ecotype inhabiting an environmentally variable bioregion had higher upper thermal tolerance and stronger plastic responses to warming than temperate and desert ecotypes (Sandoval-Castillo et al. 2020). We therefore expected the savannah ecotype, which inhabits a more environmentally variable bioregion than the rainforest ecotype, to show a greater plastic response. This expectation was supported by the nearly twofold greater number of DE genes in the savannah ecotype, together with slightly higher mean CT_MAX_ compared to the rainforest ecotype.

Ecotype-specific responses were also reflected in differences in the protein interaction network. Because fewer DE genes were induced in the rainforest ecotype, its network was smaller and had lower connectivity. Only one hub gene, *ACLYB*, was responsive in both the savannah and rainforest ecotypes. This gene supplies acetyl-CoA for multiple biochemical reactions in protein, carbohydrate and lipid metabolism, and altered expression during heat shock stress may promote osmoprotectant and antioxidant responses, thereby enhancing cellular heat tolerance (Chan et al. 2023). In contrast, eight uniquely induced genes were identified as hubs in the savannah network, all of which are involved in the sequential synthesis of cholesterol, an important component of the heat shock response (Solano et al. 2025). These differences suggest that the rainforest and savannah ecotypes engage different molecular pathways in response to warming, with the savannah response involving a broader range of molecular functions. Genome-wide diversity is similar between rainforest and savannah rainbowfish populations (Gates et al. 2026), suggesting that the more limited transcriptional response of the rainforest ecotype is unlikely to be explained simply by reduced standing genetic variation associated with its naturally fragmented habitat (Gates et al. 2023).

Instead, we hypothesize that differences in hydroclimatic regimes, landscape structure, and connectivity between bioregions shape the distinct plastic responses of rainbowfish ecotypes to climate change. Despite their geographic proximity, the savannah and rainforest bioregions differ markedly in climate and ecology (Ash 1988), as also supported by the significant niche identity test. In the Australian Wet Tropics, savannahs experience strong thermal and hydrological variability, with ephemeral lowland catchments becoming connected during high-volume tropical monsoons (Ash 1988; Bowman et al. 2010; Howley et al. 2013). In contrast, rainforests are relatively stable and aseasonal, with semi-isolated catchments embedded in rugged, complex, and fragmented terrain (Murphy and Bowman 2012; Ondei et al. 2017; Gates et al. 2023).

The potential role of connectivity and environmental heterogeneity in shaping adaptation in this system was recently examined using genotype–phenotype–environment associations in the two ecotypes (Gates et al. 2026). Across 510 rainbowfish from 17 localities, genomic and morphological variation was more strongly associated with bioclimatic heterogeneity than with neutral or spatial factors. Phenotype–environment associations were strong in both ecotypes, particularly in savannah populations. Rainforest populations showed stronger signals of local genetic adaptation to hydroclimatically stable environments, whereas the highly connected savannah populations, which experience more variable and heterogeneous conditions, appeared to rely less on polygenic adaptation and more on phenotypic plasticity to track environmental variation (Gates et al. 2026). Together with our finding that the number and identity of differentially expressed genes differ between ecotypes, these results raise the possibility that rainforest specialists have experienced stronger local adaptation and greater genetic assimilation of thermal responses than savannah generalists (Kelly 2019; Scheiner and Levis 2021). More broadly, these findings highlight a role for bioregional differences in habitat variability and landscape complexity in shaping thermal resilience and evolutionary trajectories of tropical organisms.

### Contrasting plasticity and resilience to climate change across a continent

Regional differences in climate can drive substantial geographic variation in organismal physiology, including thermal tolerance (Cohet et al. 1980; Compton et al. 2007; Sunday et al. 2019). Consequently, adaptive resilience to climate change is expected to vary geographically (Thomas et al. 2004; Román-Palacios and Wiens 2020; Trisos et al. 2020). Assuming that populations are adapted to the environments they experience, Janzen (1967) proposed that the relatively stable climates of the tropics should favour thermal specialisation, whereas greater climatic variability at higher latitudes should favour thermal generalists. This framework has since been extensively tested along latitudinal gradients, with evidence that tropical organisms generally have higher thermal maxima but narrower thermal safety margins than temperate organisms (Addo-Bediako et al. 2000; Ghalambor et al. 2006; Deutsch et al. 2008; Sunday et al. 2011; Sunday et al. 2019).

To compare patterns of climate-change resilience across Australia, we integrated and re-analysed RNA-seq data from subtropical, desert and temperate rainbowfish ecotypes (Sandoval-Castillo et al. 2020) against an improved reference transcriptome, adding new data from tropical rainforest and savannah ecotypes. A phylogenetic analysis of genome-wide data showed that the ecotypes are closely related, consistent with a broader phylogeny of the family Melanotaeniidae based on mitochondrial DNA and the nuclear S7 gene (Unmack et al. 2013). Given this recent shared ancestry, differences in physiological responses likely reflect both neutral evolutionary divergence and adaptation to contrasting bioclimatic regions (Sandoval-Castillo et al. 2020). Transcriptional responses to the simulated heat event were largely ecotype-specific: no climate-responsive genes were shared across all five ecotypes, and only six were shared by at least four ecotypes. On average, just 3.7% (range 0.8–10.6%) of the 520 climate-responsive genes were shared across the ten pairwise ecotype comparisons. Hub genes in the protein interaction network of the ecotypes were predominantly associated with heat shock protein complexes and sterol/cholesterol biosynthesis, indicating central roles in the thermal response. Enriched pathways were similarly dominated by stress-response processes, particularly heat shock and oxidative stress responses. Together, these patterns suggest that protein homeostasis, redox balance and lipid metabolism may contribute to thermal resilience. Heat shock and lipid metabolic pathways have previously been implicated in resilience and adaptive responses to rapid climatic change (Logan and Cox 2020; Zhang and Dong 2021). We therefore propose the DE genes identified here, particularly the hub genes, as promising candidates for further investigation of the molecular basis of climate adaptation in rainbowfishes.

The magnitude of these climate-induced expression responses was strongly positively associated with thermal tolerance across the five ecotypes (Figure 4). Plasticity and thermal performance were highest in the tropical-savannah and subtropical ecotypes (both from environmentally variable bioregions), intermediate in the desert ecotype, and lowest in the temperate ecotype. The tropical-rainforest ecotype was the exception, showing high thermal tolerance but limited plasticity. Consistent with this trend, positive associations between gene expression plasticity and ecologically adaptive thermal tolerance have been reported in teleosts (*Oncorhynchus mykiss gairdneri* ecotypes, Narum and Campbell 2015; *Neogobius melanostomus* versus *Proterorhinus semilunaris*, Wellband and Heath 2017; *Menidia beryllina* versus *Hypomesus transpacificus*, Komoroske et al. 2021) and oysters (*Crassostrea angulata* versus *Crassostrea gigas*, Li et al. 2021). Transcriptional flexibility may therefore be an important determinant of physiological stress resilience (Ghalambor et al. 2015; Bay et al. 2017; Kelly 2019), and trait plasticity is increasingly recognised to evolve via ecological selection as a phenotypic trait in its own right (Scheiner 1993; Pigliucci 2001; Chevin and Lande 2015; Bailey et al. 2021; Bond et al. 2021). Although the rainbowfish patterns do not demonstrate causation, they support the hypothesis that bioregional climatic history shapes broad patterns of climate resilience and adaptation, including transcriptional plasticity and thermal tolerance.

The average CT_MAX_ of the tropical and subtropical ecotypes was significantly higher than that of the other rainbowfishes, with thermal tolerance generally decreasing with distance from the equator. These two ecotypes also had the highest environmental warming tolerance. These patterns are unlikely to be explained by phylogeny alone. Although the two tropical ecotypes (*M. s. splendida*) and the desert ecotype (*M. s. tatei*) belong to the same species, the desert ecotype had comparatively low CT_MAX_ and environmental warming tolerance. Similarly, the warm-tolerant subtropical ecotype (*M. duboulayi*) is closely related to the temperate *M. fluviatilis* (Figure 1B; Unmack et al. 2013), which had the lowest CT_MAX_ and environmental warming tolerance of all ecotypes. Together, these comparisons are consistent with climatic adaptation. However, as with gene-expression plasticity, interpreting these patterns may require consideration of both average climatic conditions and climatic variability. A survey of the aquatic literature suggests that species inhabiting environments characterised by highly variable thermal extremes induce the heat shock response at maximal or near-maximal levels (Tomanek 2010). Because heat shock response activity indicates thermal stress and proximity to tolerance limits, such species may have less acclimatory capacity and greater vulnerability to climate change (Tomanek 2010). This may help explain the unexpectedly low climatic resilience of the temperate and desert ecotypes, which inhabit highly variable and extreme environments. The difference between the mean coldest and warmest monthly temperatures across their distributions is 27.9°C and 30.8°C, respectively — the highest of all ecotypes (Table S7). Across their ranges, both ecotypes also show strong genotype–environment associations with hydroclimatic variation. In *M. fluviatilis*, adaptive genetic diversity is associated with seasonal variation in stream flow and the timing of temperature and precipitation extremes in temperate and semi-arid environments (Brauer et al. 2018). In *M. s. tatei*, adaptive genetic diversity correlates with aridity, particularly in extremely unstable rivers and waterholes of arid central-western Australia (Attard et al. 2022). Large water bodies can buffer extreme air temperatures, but climate change is expected to intensify heatwaves and droughts, reducing aquatic refugia in arid and semi-arid regions (Magoulick and Kobza 2003) and increasing fishes’ exposure to heat stress. As refugia contract, behavioural responses such as moving to deeper or cooler water may also become less effective. Understanding how climate change alters the microenvironmental features that support behavioural thermoregulation (Magoulick and Kobza 2003; Kearney et al. 2009) will therefore be important for assessing vulnerability in desert aquatic ecosystems.

Applying the same rationale to the tropical ecotypes, the greater climatic variability of the savannah could impose a trade-off for maximum thermal tolerance relative to the more stable rainforest. However, although warm thermal extremes in the savannah are substantially higher (∼2.2°C), cold extremes are only slightly lower (∼0.4°C), potentially reducing selection for tolerance of cold conditions. Savannah organisms may therefore benefit more consistently from adaptation to warm conditions than the rainforest ecotype, as reflected in their greater thermal tolerance and associated expression plasticity. Flexibility in thermal responses may be particularly important for persistence under warmer and more variable future climates (Logan and Buckley 2015; Diamond and Martin 2021), suggesting that the savannah ecotype may possess the greatest preadapted resilience among the ecotypes studied here. The rainforest ecotype also has relatively high thermal tolerance, but evidence of climatic specialisation suggests that its responses may be less flexible outside the environmental conditions to which it is adapted.

## Conclusions and directions

Our results highlight strong differences in plastic capacity and thermal tolerance among closely related rainbowfishes, cautioning against treating ecotypes and species as homogeneous units in distribution forecasts (Pearman et al. 2010; Reed et al. 2011; King et al. 2019). Realised vulnerability will depend not only on existing adaptations but also on the magnitude and nature of local environmental change (Huntley et al. 1995; Sykes et al. 1996). Future studies should therefore incorporate these differences into distribution models under projected climates. However, predicting future adaptive potential from contemporary patterns remains challenging, because responses to acute thermal stress may differ from those favoured by warmer long-term conditions (Gerken et al. 2015). Heatwaves may impose particularly strong selection, and heat shock and lipid metabolic pathways are plausible targets under rapid climatic change (Logan and Cox 2020; Zhang and Dong 2021). The DE genes identified here are candidates for adaptation to future climates, particularly the hub genes occupying central positions in the expression network. Testing for divergent selection in the evolution of expression variation in candidate genes (Rohlfs and Nielsen 2015) across hydroclimatic and latitudinal gradients could help identify ecotype-specific responses to climate change (e.g. Brauer et al. 2017; Sandoval-Castillo et al. 2020).

Linking transcriptomic and fitness experiments, mechanistic niche models, landscape genomics, and genetic architecture will be important for predicting evolutionary responses and mitigating biodiversity loss under future climates (Bay et al. 2017; Grummer et al. 2019; Bernatchez et al. 2024). Broader studies across co-distributed taxa and assessments of thermal tolerance under both laboratory and natural conditions will further clarify the role of plasticity in climate responses. Nonetheless, the strong biogeographic patterns in warming tolerance and expression plasticity observed across Australian rainbowfishes provide evidence for ecological differentiation in thermal plasticity. Predicting winners and losers of climate change remains challenging, but our results suggest that biogeographic variation in plasticity may be an important determinant of population persistence and species distributions under global warming.

## Author Contributions

L.B.B. and L.B. conceived the study with assistance from J.S.-C. and C.J.B. L.B.B. and L.B. secured funding and resources. J.S.-C., K.G., and C.J.B. generated the data. All authors collected samples. J.S.-C., K.G. and C.J.B. performed data analysis. All authors contributed to the interpretation of results. L.B.B. supervised the research. L.B.B. and K. G. drafted the manuscript; coauthors revised the manuscript and approved its final version.

## Acknowledgements

We thank Leslie Morrison and her exceptional team at Flinders University Biodiversity Facility for the fish husbandry over the years, Michael Hammer and Keith Martin for sharing their knowledge of rainbowfish ecology and distributions, Leo O’Reilly, Catherine Attard, Steve Smith and David Schmarr for assistance with sampling, and Diana-Elena Vornicu for preparing the rainbowfish illustrations, which were based on photographs provided by the authors and by Gunther Schmida. We acknowledge the Traditional Owners of the several regions on which this research was conducted, including the Eastern Kuku Yalanji people of the Mossman and Daintree regions, the Ang-Gnarra people of the Normanby catchment, the Turrbal and Jagera (Yuggera) peoples of the Brisbane River, the Arabana people of the Lake Eyre and the Neales River system, and the Barkindji (Paakantyi) and Latji Latji peoples of the middle Murray River, and pay our respects to Elders past and present. We are also grateful to colleagues from MELFU and CEBEL at Flinders University, as well as the Labo Bernatchez at Laval University, for their insightful discussions and assistance with methodological approaches. Field collections were performed under permits from various government agencies from South Australia, New South Wales and Queensland. Our rainbowfish research has been funded by the Australian Research Council (DP110101207, DP150102903, DP190102533, FT130101068, and DP260104361 to LBB). Additional support was received by the Royal Society of South Australia for the Advancement of Science, and the Flinders University Student Association Development Grant. KG received support from the AJ & IM Naylon PhD Scholarship through Flinders University and the Playford Trust/Thyne Reid Foundation PhD Scholarship.

## Data Accessibility Statement

The data have been archived at figshare: https://figshare.com/s/9a738ed6eb19575a9508

10.6084/m9.figshare.34031337

## Ethical Approval

Sampling, transport and animal handling procedures were performed in accordance with Flinders University Animal Ethics Committee approvals E342, E429, and E463.

## Benefits Generated

Benefits from this research accrue from the sharing of our data and results on public databases as described above.

