## Supplemental material for "From tropical rainforests to scorching deserts: biogeography shapes climate-change plasticity in Australian rainbowfishes"

### Table of Contents:

| **Figure S1** | Page 2 |
| --- | --- |
| **Figure S2** | Pages 3 and 4 |
| **Table S1** | Page 5 |
| **Table S2** | Page 5 |
| **Table S3** | Page 6 |
| **Table S4** | Pages 7 and 8 |
| **Table S5** | Page 9 |
| **Table S6** | Page 9 |
| **Table S7** | Page 10 |
| **Table S8** | Extended Table_S8.xlsx |

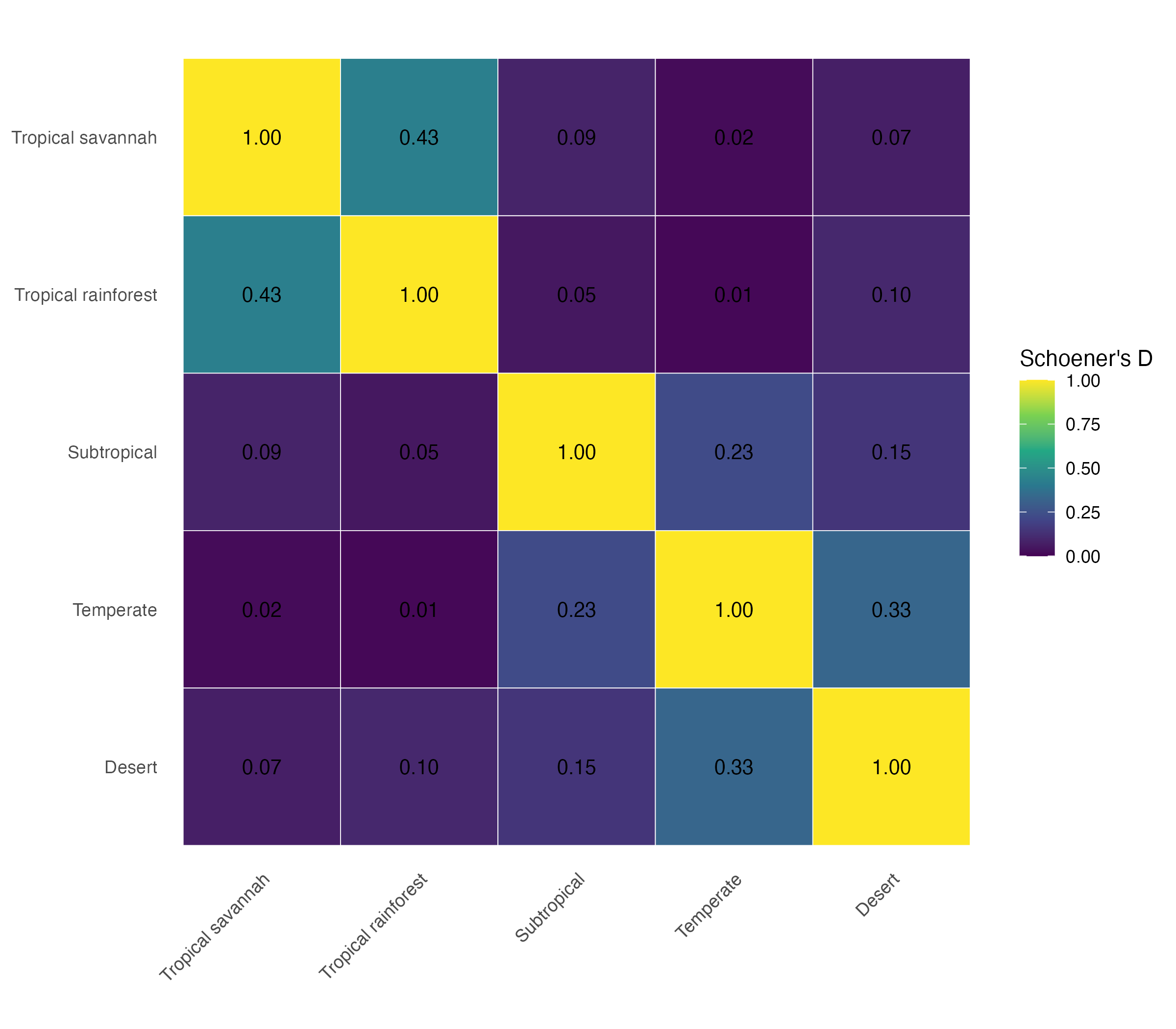

**Figure S1.** Ecotype niche overlap for the five *Melanotaenia* ecotypes.

A

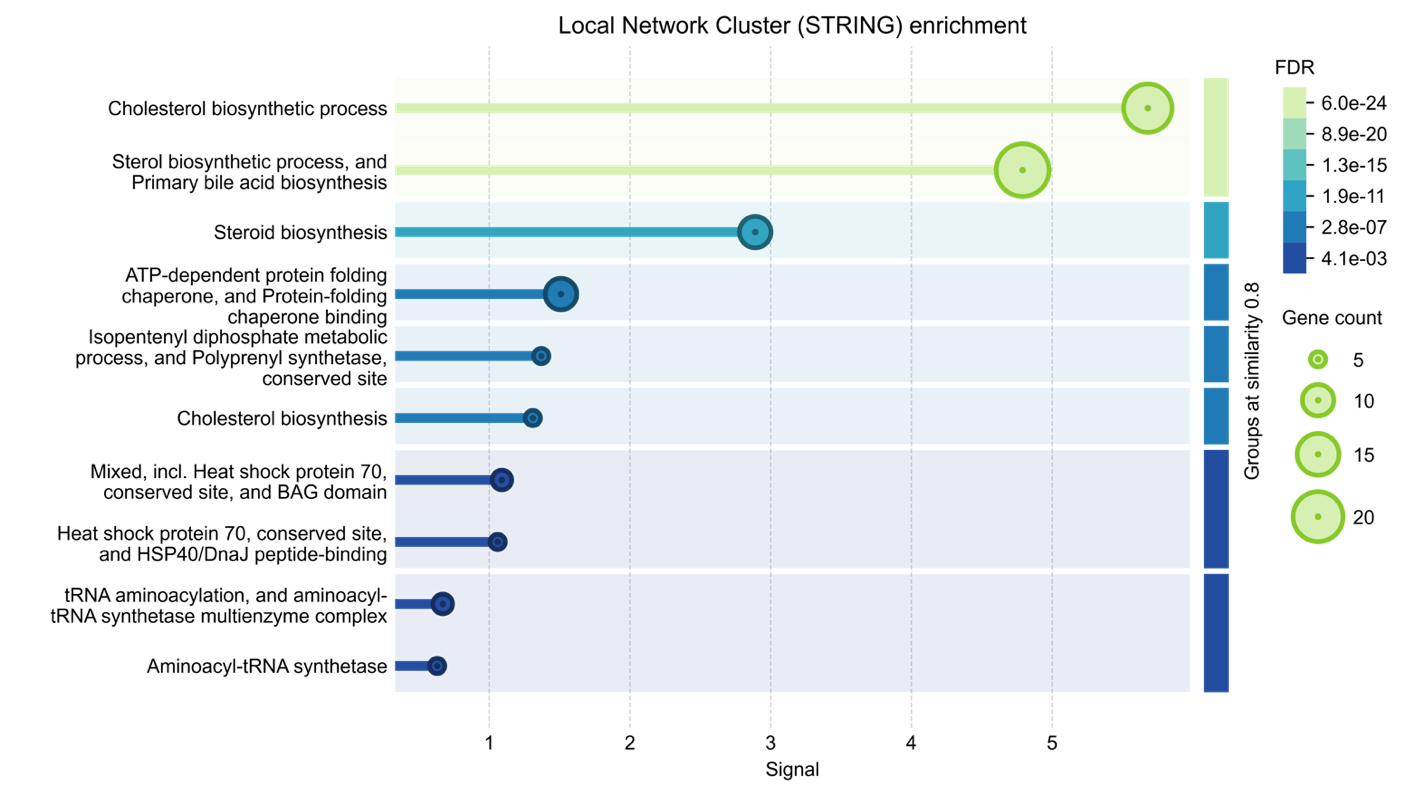

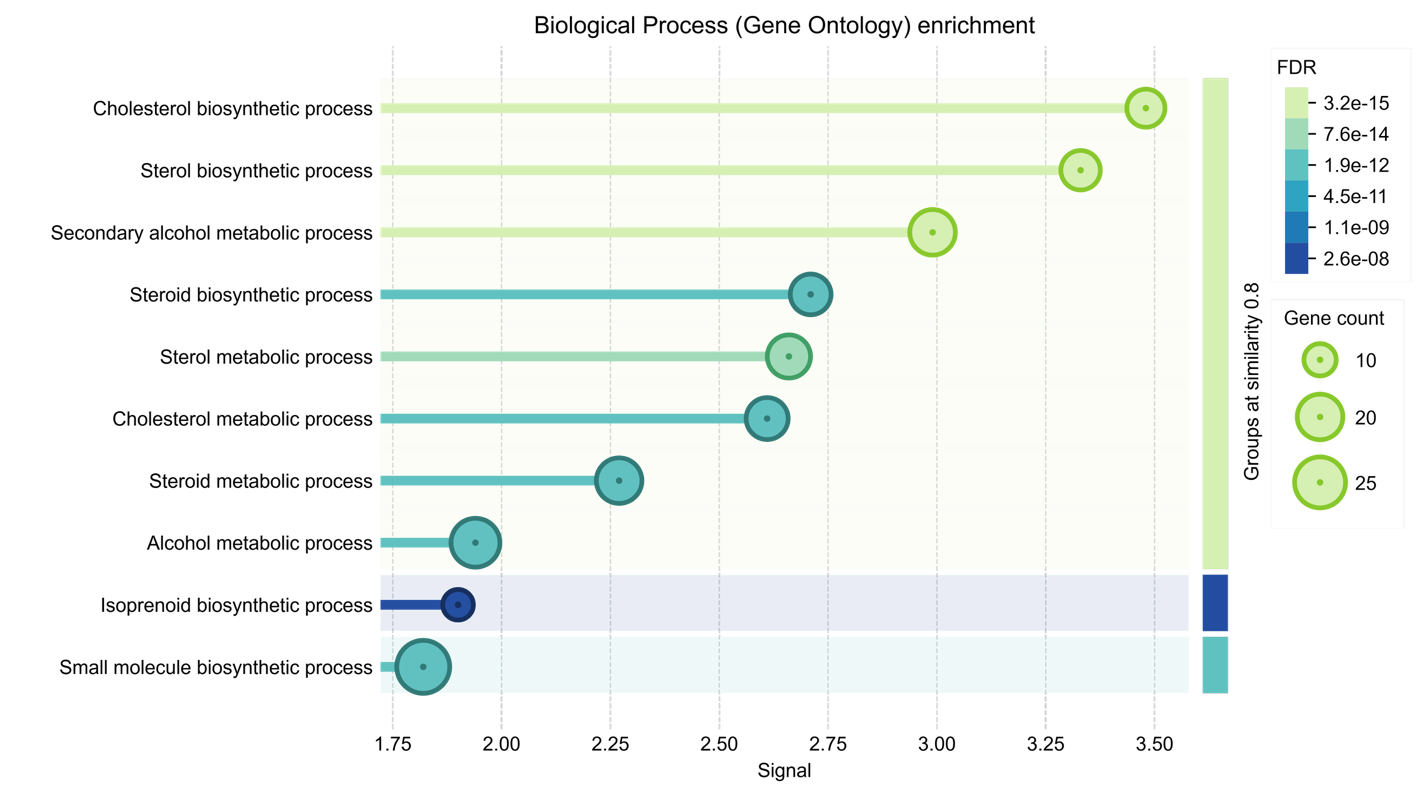
 **Figure S2.** Functional enrichment among thermally differentially expressed genes, from STRING and Gene Ontology analyses. **(A)** STRING local network cluster enrichment. **(B)** Gene Ontology Biological Process enrichment. **(C)** Gene Ontology Cellular Component enrichment. **(D)** Gene Ontology Molecular Function enrichment. In all panels, terms with ≥ 0.8 similarity are grouped (right-hand colour bar); within each term, dot size reflects gene count and dot/line colour reflects FDR-corrected significance. The x-axis (signal) reflects a composite enrichment score combining term specificity and significance. Note that the gene-count scale is set independently for each panel and is not directly comparable across panels.

B

C

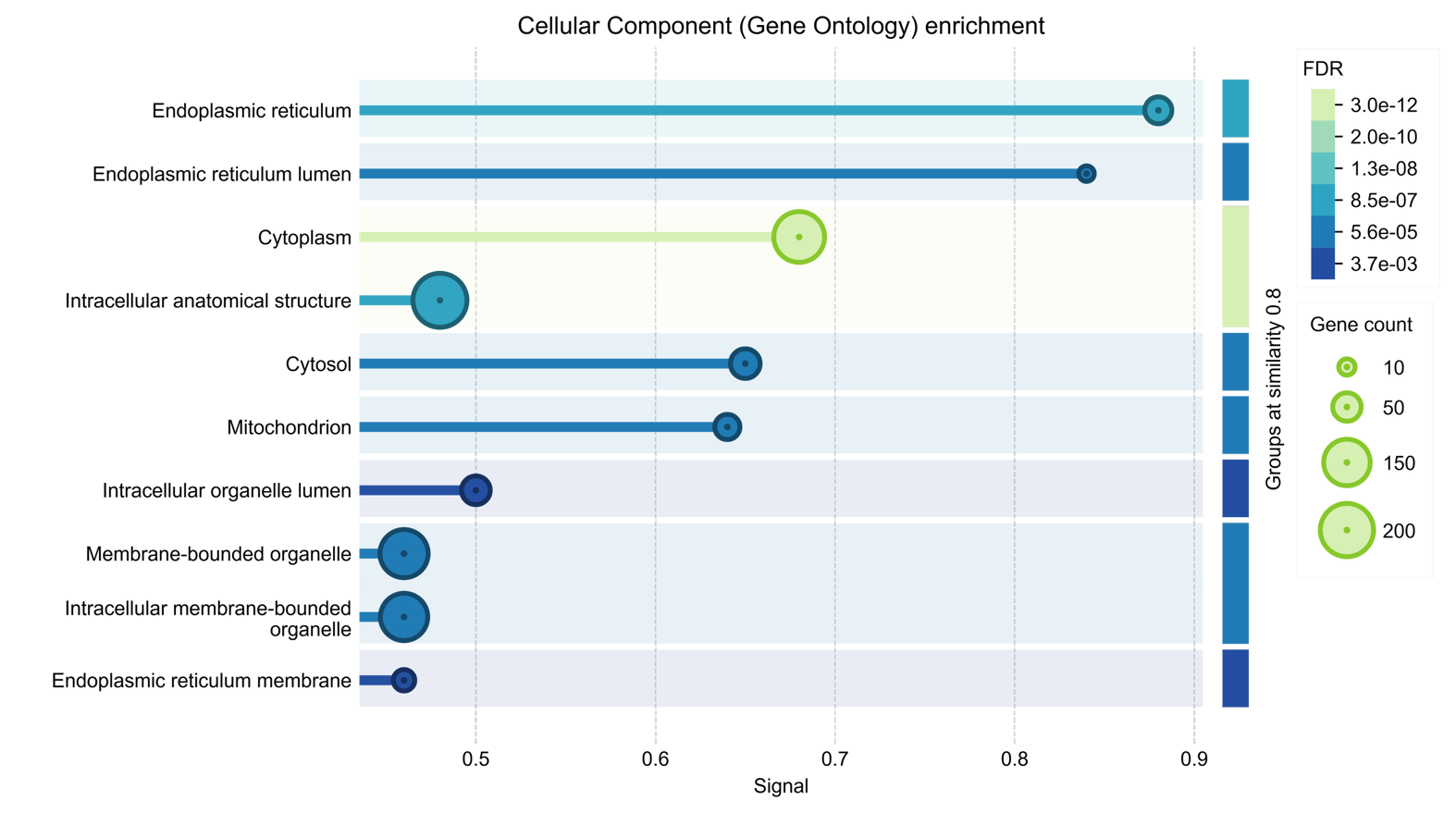

D

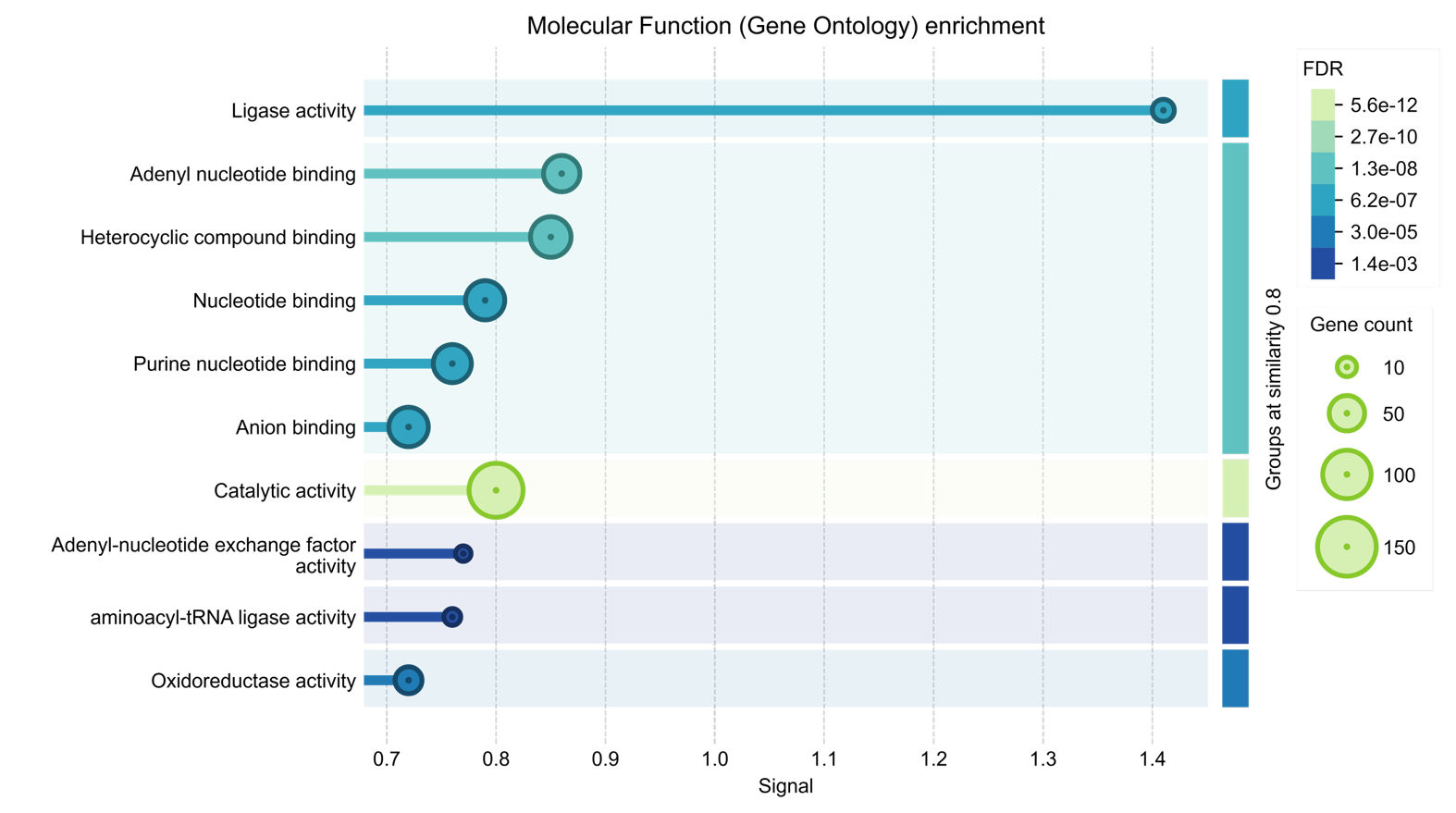

**Figure S2.** (Continued)

**Table S1**. Niche model support for the five *Melanotaenia* ecotypes and variable predictors

| **Ecotype** | **Occurrences** | **Background** | **Predictors** | **auc** | **auc_sd** | **Omission** |
| --- | --- | --- | --- | --- | --- | --- |
| Savannah | 188 | 2958 | bio5;bio6;bio12;bio15;bio1;bio14;bio3 | 0.842 | 0.102 | 0.069 |
| Rainforest | 87 | 1391 | bio5;bio6;bio12;bio15;bio2;bio4;bio1;bio3 | 0.803 | 0.119 | 0.125 |
| Subtropical | 446 | 2953 | bio5;bio6;bio12;bio15;bio2;bio3 | 0.609 | 0.069 | 0.095 |
| Temperate | 557 | 3000 | bio5;bio6;bio12;bio15;bio2;bio18;bio9;bio8;bio3 | 0.621 | 0.123 | 0.245 |
| Desert | 210 | 3000 | bio5;bio6;bio12;bio2;bio14;bio9;bio8;bio3 | 0.537 | 0.102 | 0.144 |

**Table S2**. Niche identity tests for the five *Melanotaenia* ecotypes

| Ecotype 1 | Ecotype 2 | D | I | p_D | p_I | permutations | p_D_holm | p_I_holm |
| --- | --- | --- | --- | --- | --- | --- | --- | --- |
| Savannah | Rainforest | 0.430 | 0.736 | 0.001 | 0.001 | 999 | 0.01 | 0.01 |
| Savannah | Subtropical | 0.085 | 0.252 | 0.001 | 0.001 | 999 | 0.01 | 0.01 |
| Savannah | Temperate | 0.022 | 0.112 | 0.001 | 0.001 | 999 | 0.01 | 0.01 |
| Savannah | Desert | 0.072 | 0.229 | 0.001 | 0.001 | 999 | 0.01 | 0.01 |
| Rainforest | Subtropical | 0.049 | 0.193 | 0.001 | 0.001 | 999 | 0.01 | 0.01 |
| Rainforest | Temperate | 0.014 | 0.113 | 0.001 | 0.001 | 999 | 0.01 | 0.01 |
| Rainforest | Desert | 0.098 | 0.285 | 0.001 | 0.001 | 999 | 0.01 | 0.01 |
| Subtropical | Temperate | 0.227 | 0.470 | 0.001 | 0.001 | 999 | 0.01 | 0.01 |
| Subtropical | Desert | 0.147 | 0.378 | 0.001 | 0.001 | 999 | 0.01 | 0.01 |
| Temperate | Desert | 0.330 | 0.632 | 0.001 | 0.001 | 999 | 0.01 | 0.01 |

| **Table S3.** Summary of assembly statistics for the *Melanotaenia* transcriptome. ORF = open reading frame, bp = base pairs \|; Raw read pairs \| 1,829,323,181; Retained filtered read pairs \| 1,144,838,067; Transcripts \| 450,246; Trinity Genes \| 315,268; Trinity Genes with ORF \| 64,694; Uni-genes \| 36,536; N50 \| 1,497; Average contig length (bp) \| 1,070; Assembled bases \| 39,247,621; ORF genes annotated \| 26,919 **Table S4.** Number (n) and percentages (%) of all characterised genes of *Melanotaenia* ecotypes and the corresponding differentially expressed genes between treatment groups | | |
| --- | --- | --- |
| **Comparisons based on 28,181 sequenced genes** | **%** | **n** |
| TemperateRainforest | 89.7 | 32,772 |
| TemperateDesert | 89.4 | 32,677 |
| TemperateSubtropical | 88.8 | 32,427 |
| TemperateSavannah | 90.1 | 32,903 |
| RainforestDesert | 87.8 | 32,061 |
| RainforestSubtropical | 86.0 | 31,413 |
| RainforestSavannah | 88.3 | 32,256 |
| DesertSubtropical | 85.9 | 31,388 |
| DesertSavannah | 88.3 | 32,258 |
| SubtropicalSavannah | 86.4 | 31,583 |
| TemperateRainforestDesert | 85.0 | 31,070 |
| TemperateRainforestSubtropical | 84.1 | 30,743 |
| TemperateRainforestSavannah | 85.5 | 31,245 |
| TemperateDesertSubtropical | 84.2 | 30,751 |
| TemperateDesertSavannah | 85.6 | 31,267 |
| TemperateSubtropicalSavannah | 84.6 | 30,915 |
| RainforestDesertSubtropical | 82.2 | 30,036 |
| RainforestDesertSavannah | 84.4 | 30,819 |
| RainforestSubtropicalSavannah | 82.5 | 30,157 |
| DesertSubtropicalSavannah | 82.8 | 30,238 |
| TemperateRainforestDesertSubtropical | 80.8 | 29,529 |
| TemperateRainforestDesertSavannah | 82.2 | 30,024 |
| TemperateRainforestSubtropicalSavannah | 81.1 | 29,639 |
| TemperateDesertSubtropicalSavannah | 81.4 | 29,730 |
| RainforestDesertSubtropicalSavannah | 79.9 | 29,181 |
| TemperateRainforestDesertSubtropicalSavannah | 78.8 | 28,781 |
| **Comparisons based on 520 differently expressed genes** | **%** | **n** |
| TemperateRainforest | 1.5 | 8 |
| TemperateDesert | 2.5 | 13 |
| TemperateSubtropical | 1.7 | 9 |
| TemperateSavannah | 0.8 | 4 |
| RainforestDesert | 6.3 | 33 |
| RainforestSubtropical | 1.7 | 9 |
| RainforestSavannah | 10.6 | 55 |
| DesertSubtropical | 2.7 | 14 |
| DesertSavannah | 6.7 | 35 |
| SubtropicalSavannah | 2.5 | 13 |
| TemperateRainforestDesert | 0.8 | 4 |
| TemperateRainforestSubtropical | 0.2 | 1 |
| TemperateRainforestSavannah | 0.4 | 2 |
| TemperateDesertSubtropical | 0.8 | 4 |
| TemperateDesertSavannah | 0.8 | 4 |
| TemperateSubtropicalSavannah | 0.2 | 1 |
| RainforestDesertSubtropical | 1.0 | 5 |
| RainforestDesertSavannah | 4.6 | 24 |
| RainforestSubtropicalSavannah | 0.8 | 4 |
| DesertSubtropicalSavannah | 0.8 | 4 |
| TemperateRainforestDesertSubtropical | 0.0 | 0 |
| TemperateRainforestDesertSavannah | 0.4 | 2 |
| TemperateRainforestSubtropicalSavannah | 0.0 | 0 |
| TemperateDesertSubtropicalSavannah | 0.2 | 1 |
| RainforestDesertSubtropicalSavannah | 0.6 | 3 |
| TemperateRainforestDesertSubtropicalSavannah | 0.0 | 0 |

**Table S5.** Thermal tolerance comparisons based on critical thermal maximum (CT_MAX_) of five rainbowfish ecotypes with a single factor ANOVA.

| **Species** | **Ecotype** | ***n*** | **Max CT_MAX_** | **Min CT_MAX_** | **Average** | **Variance** |
| --- | --- | --- | --- | --- | --- | --- |
| *Melanotaenia splendida splendida* | Tropical savannah | 10 | 39.0 | 37.9 | 38.40 | 0.129 |
| *Melanotaenia splendida splendida* | Tropical rainforest | 10 | 38.9 | 37.5 | 38.23 | 0.293 |
| *Melanotaenia duboulayi* | Subtropical | 10 | 38.6 | 37.0 | 37.96 | 0.174 |
| *Melanotaenia splendida tatei* | Desert | 10 | 37.6 | 36.1 | 37.24 | 0.232 |
| *Melanotaenia fluviatilis* | Temperate | 10 | 36.5 | 33.1 | 34.88 | 1.633 |

| **Source of Variation** | **SS** | **df** | **MS** | **F** | **P-value** | **F crit** |
| --- | --- | --- | --- | --- | --- | --- |
| Between Groups | 83.6168 | 4 | 20.9042 | 42.4786182 | 9.9E-15 | 2.578739184 |
| Within Groups | 22.145 | 45 | 0.49211111 |  |  |  |
| Total | 105.7618 | 49 |  |  |  |  |

**Table S6.** Thermal tolerance comparisons based on critical thermal maximum (CT_MAX_) of five rainbowfish ecotypes using two-sample *t*-tests.

|  | **Rainforest vs Savannah** | | **Tropical vs Subtropical** | | **Tropical vs Desert** | | **Tropical vs Temperate** | |
| --- | --- | --- | --- | --- | --- | --- | --- | --- |
|  | *Variable 1* | *Variable 2* | *Variable 1* | *Variable 2* | *Variable 1* | *Variable 2* | *Variable 1* | *Variable 2* |
| Mean | 38.230 | 38.400 | 38.315 | 37.960 | 38.315 | 37.240 | 34.880 | 38.315 |
| Variance | 0.293 | 0.129 | 0.208 | 0.174 | 0.208 | 0.232 | 1.633 | 0.208 |
| Observations | 10.000 | 10.000 | 20.000 | 10.000 | 20.000 | 10.000 | 10.000 | 20.000 |
| Pooled Variance | 0.211 |  | 0.197 |  | 0.215 |  |  |  |
| Hypothesized Mean Difference | 0.000 |  | 0.000 |  | 0.000 |  | 0.000 |  |
| *df* | 18.000 |  | 28.000 |  | 28.000 |  | 10.000 |  |
| *t* Stat | -0.827 |  | 2.066 |  | 5.981 |  | -8.243 |  |
| *P*(*T*<=*t*) one-tail | 0.209 |  | 0.024 |  | 0.000 |  | 0.000 |  |
| *t* Critical one-tail | 1.734 |  | 1.701 |  | 1.701 |  | 1.812 |  |
| *P*(*T*<=*t*) two-tail | 0.419 |  | 0.048 |  | 0.000 |  | 0.000 |  |
| *t* Critical two-tail | 2.101 |  | 2.048 |  | 2.048 |  | 2.228 |  |

| **Table S7** Temperature range across the distribution of the *Melanotaenia* ecotypes based on baseline WorldClim 2.1 bioclimatic variables (1970–2000) | | | | | |
| --- | --- | --- | --- | --- | --- |
| **Ecotype** | **Period** | **Number of occurrence cells** | **Mean coldest month** | **Mean Warmest Month** | **Temperature range** |
| Tropical savannah | 1970-2000 | 188 | 14.3 | 32.1 | 17.8 |
| Tropical rainforest | 1970-2000 | 87 | 15.2 | 30.6 | 15.4 |
| Subtropical | 1970-2000 | 446 | 7.6 | 29.1 | 21.5 |
| Temperate | 1970-2000 | 557 | 4.3 | 32.2 | 27.9 |
| Desert | 1970-2000 | 210 | 6.7 | 37.5 | 30.8 |
